# Artificial Intelligence for Bioinformatics: Applications in Protein Folding Prediction

**DOI:** 10.1101/561027

**Authors:** Max Staples, Leong Chan, Dong Si, Kasey Johnson, Connor Whyte, Renzhi Cao

## Abstract

AI recently shows great promise in the field of bioinformatics, such as protein structure prediction. The Critical Assessment of protein Structure Prediction (CASP) is a nationwide experiment that takes place biannually, which centered around analyzing the best current systems for predicting protein tertiary structures. In this paper, we research on available AI methods and features, and then explore novel methods based on reinforcement learning. Such method will have profound implications for R&D in bioinformatics and add an additional platform to the management of innovation in biotechnology.

## I. Introduction

Bioinformatics is an extremely popular field in the development of artificial intelligence[1]–[7]. It shows great promise into applications such as gene therapy, personalized medicine, preventative medicine, drug development, and biotechnology [6]. In order for many of these applications to exist, the native structure of the protein must be known. However, it is still difficult to find the structure of a protein.

The most effective methods for finding protein structure are time consuming and expensive. These methods are Nuclear Magnetic Resonance and X-ray crystallography. More recently, cryo-electron microscope (cryo-EM) as a revolutionary technique has been used to efficiently produce high-resolution large-scale molecular structures. Machine learning and artificial intelligence have also been utilized on decipher these cryo-EM density maps [7]–[10]. Perhaps the most complex part of obtaining these scans is getting the protein to crystalize in order to run these tests. Often many liquid proteins with not even crystalize. Artificial intelligence is a possible better pathway to sequencing these proteins.

Artificial Intelligence is promising in the field[11], [12] because it has been successfully applied in different fields like business for identifying job-hopping patterns[13], image recognition, etc. It can accurately and efficiently predict thousands of possible structures. The issue is most of the models are inaccurate and do not produce predicted proteins that contain useful information. Using artificial intelligence, programs are trained using many numerically represented atomic features from the models (such as bond lengths, bond angles, residue-residue interactions, physio-chemical properties, and potential energy properties) to compare the prediction models output to the known crystal structures to assess the quality of the model and find the most accurate model. Models for predictions and prediction analysis are compared each year in one main gathering called the Critical Assessment of Structure Prediction (CASP), which is the flagship experiment of protein structure prediction. Every two years researchers from around the world submit machine learning methods designed for protein structure prediction[14], especially break-through method MULTICOM in the recent CASP 13, while deep learning has been applied to make protein structure prediction with help of protein contact distance prediction[15]. These methods are analyzed for performance by professionals in the field. Neither the participants nor the assessors are aware of one another’s identity ensuring the quality of the experiment. Over time the CASP experiment has evolved and the categories within it have changed significantly. One important addition to the CASP was the quality assessment (QA) category added in CASP7 (2006) to analyze the performance of the models given the broad range of accuracy that predicted structures span.[1].

## II. Background

In this section, we do a literature review on the current research in the field of protein structure prediction and the quality assessment of predicted structures. We begin with a review of the current state of the CASP experiment with special emphasis on the QA category. Next, we present a review of a number of methods currently dominating the field. For each method we have determined what makes the method unique as well as its pros and cons.

### A. CASP Experiment XII[1]

The CASP recently held its 13 th bi-annual experiment, however, the analysed results have not been released so we focus on the state of the CASP 12. In this CASP, there were 188 methods submitted from 96 research groups in 19 different countries. In total there were 55,000 predictions, 7400 of which were QA predictions.

The QA category can be broken into two subcategories – multi-model and single-model QA methods[16]. Multi-model methods rely on a group of models predicted for one target and determine poor models, in part, by searching for outlier models within the group. This leads to poor selection of the best overall model[17]. Single-model methods focus on predicting accuracy for one predicted structure. Historically, consensus methods have performed better than single-model methods, however, that has changed as of CASP 12. The single-model methods now have lower error with respect to predicted GDT_TS scores compared to true quality scores. This gain has been attributed to improvements in energy functions and better machine learning strategies.

### B. Neural Networks

Neural networks are useful tools in solving protein prediction problems because a neural network can assign various weights to features depending on learned probabilities from a dataset. The network is able to see trends in how proteins fold and then based on assigned weight predict how a new protein would fold. Neural networks have the ability to predict things like contact areas and folding energy in order to better predict a tertiary protein structure. Used in tandem with multiple physio-chemical properties, neural networks are a highly accurate method.

Here we list three state-of-the-art QA tools:

- *ProQ3D[18].* Proq has been a top performing method since its original release in 2003. Since that time the method has gone through several iterations. The original method relied on a neural network and extensive features. The next two iterations (proq2, proq3) relied on a support vector machine and additional features. Proq2 received profile weights as new feature and saw an improvement in prediction as a result, and in Proq3 an energy function was added also leading to an improvement in prediction. Given the recent success of deep learning, the authors decided to implement a deep neural network on proq3 and proq2. Doing this resulted in an increase in global correlation from 0.85 to 0.90 for proQ3D and 0.81 to 0.85 for proQ2D.
- *QACon[19].* QAcon used 12 features to train AI model for QA problem. These features utilize structural features, physiochemical properties, residue contact predictions. The residue contact prediction methods used are PSICOV and DNcon. The training technique utilized is a neural network trained using the 12 features with two layers. The method was trained using datasets provided in CASP 9, and tested on datasets from CASP 11. The method tested as one of the top single-model QA. The method was tested without the added residue-residue contact and performed much worse than the original. This showed the importance and impact fullness of the added feature.
- *DeepQA[20].* This paper introduces another novel single-model quality assessment. This method utilizes 16 features including 9 top-performing energy and knowledge-based potential scores and seven based on physiochemical properties. The deep belief network is trained utilizing data sets provided from previous CASP experiments. They also trained using several publicly available data sets and models generated by in house ab-initio. The novel aspect of this method is a two layer Restricted Bultzman Machines that forms the hidden layers of the neural network. The initial weights of the RBMs are initialized using a pretraining that is unsupervised learning. It utilizes a contrastive divergence algorithm. The final step is optimized based on Broyden-Fletcher-Goldfarh-Shanno (BFGS) optimization. The training data is divided into five sets and a five-fold cross validation is used to train and validate DeepQA. Logistic regression then produces a real output of a value between 1 and 0. This deep belief network outperformed Support Vector Machine (SVM) and neural networks. It also performed well in model pools with very poor quality models, which is very important.

### C. Convolutional Neural Networks (CNNs)

CNNs take 3D representations of proteins and analyze them like images by taking high-level features such as atomic density and contact area distance from the image and compares the values of these predicted images to the actual known image of the protein structure. They utilize a similar connected structure to neural networks but are comprised of more layers that analyze different aspects of different pieces of the image. Derevyanko et al. introduced a novel quality assessment method based on CNNs that utilizes convolutional neural networks and 3D images of the predicted protein to make analysis[21]. This method is unique because it takes an image of the protein then measures the respective densities of the atoms present. It then compares this to an analysis of the native structure atomic densities and produces a score that is focused of the loss in performance of the GDT score rather than the straight GDT. Most of the analysis methods use a straight GDT score to analyze each predicted structure. Pagès et al. introduced a new tool ORNATE which relied on a 3D CNN and a local residue-wise scoring metric (CAD). The authors chose to use a local scoring metric in order to allow the network to learn locally preferred or poor 3D geometries. Each training step looked at a single residue and its local neighborhood within the structure. Some things to note were the way they chose to orient the input. Historically, a problem with 3D CNN’s has been choosing a reference orientation for the predicted structure. The authors of ORNATE decided to keep their orientation constant feeding in a cubic volumetric map of each residue and its neighborhood with the basis vectors set according to the current carbon alpha and the direction of its nitrogen bond (one basis vector determined this way, and the others follow from there). This allowed the CNN to train faster and learn more efficiently without any rotating or translating of the input. Given that many methods in the field are optimized to predict GDT_TS scores two different performance benchmarks were used for evaluation of the method. The first compared top methods in the field (SBROD, VoroMQA, RWPlus, 3dCNN, Proq3D) after training all methods on the CAD score. In this comparison, ORNATE was second to Proq3D. The second test trained all models on the GDT_TS. In this test, ORNATE performed considerably worse with respect to correlation between the predicted and true quality score, but was still the second best with respect to selecting the best overall model.

### D. Support Vector Machine (SVM)

Support Vector Machines (SVM) are supervised learning methods with associated learning methods that analyze data for regression and classification. The model represents data as points in a category which it then tries to create the longest distance between those points. SVM is widely used in bioinformatics field[4], [22]–[25], and has shown great promise and is very good at selecting models from a pool with mostly bad models. SVMQA[26] is one of the best support-vector-machine based single-model quality assessment method. This method produces a TM_Score and GDT_TS score using 19 features with 8 being potential-energy based, and 11 comparing predicted and actual models. Based on these features, a 10-fold cross-validation was performed on the training data. For each cross validation 500 trees were produced with a number of nodes, each chosen randomly. The Feature Importance Score (FIS) was then developed. The method was trained using data provided by CASP 8-10 experiments and was tested using various benchmarking datasets. The SVMQA method outperformed the best current single-model QA method in terms of both ranking structures and selecting the best model from a pool. It was also one of the best method in choosing a good model from decoys in terms of GDT loss.

### E. Probabilistic method

The probabilistic methods use the probabilities of proteins in making secondary structures and analyzing their conformation energies at each step to reevaluate the prediction it will make. After predicting secondary structures, the methods then predict a tertiary structure based on the probabilities of the secondary structures, sampling of the tertiary conformation energies, and the chemical backbone structure of the protein. UniCon3D[27] is one of the best *de novo* protein structure prediction method that performs with the top quality methods for predicting proteins while also producing lower energy conformers at each step of prediction. This model utilizes not only the structure of the protein backbone in predicting contact areas but also uses the bias of the side chains attached to the backbone. This simultaneous examination of backbone and side-chain paired with the stepwise sampling method developed (instead of what appears to be the field standard of random sampling) results in a model that is able to discern difficult targets from CASP10 and CASP11 at the same level as the top five methods with lower energy conformers which is indicative of mores stable and accurate protein structures. CONFOLD[28] is another top-performing protein structure prediction method that uses contact areas and secondary structures to better predict the final tertiary structure of the protein. It uses a unique two-stage approach that I have not seen in other papers. The first step uses contact areas and secondary structures and turns them into distances, bond angles, and hydrogen steric effects. Using these, the method looks at the predicted secondary structures that were produced and then tapers the variables that are used for the tertiary structure to patterns it detects in the initially predicted structures. This results in better resolution of beta-sheet predictions especially. This shows that weighting of the contact residue values and the number of contact areas predicted have large effect on the predicted overall structure. VoroMQA[29] is a quality assessment method that uses a unique method called Vorornoi tessellation to analyze the protein. The tessellation treats each atom as a ball and each different atom type is assigned a different sized ball. It then constructs triangles, with respective size, to represent each atom so that there are Voronoi faces where atoms are bonded. This allows for highly accurate determinations of contact areas and atom densities in a protein. This method uses a contact area distance (CAD) evaluation method where it measure the distance between contact areas (new Voronoi faces that are formed by the prediction of the tertiary structure) of the native structure and compares them to to the predicted structure. This method has shown be produce better local representations around contact areas of proteins but is slightly less effective on a global level compared to ProQ3D. Qprob[30] is a single model quality assessment method that was designed using a probability based approach. First the absolute error between a model’s feature value and the true GDT_TS score is calculated for 11 different features. Next the authors determine weights for each feature in determining an overall probability density function which is used to estimate a global quality score for a given model. This method performed in the 3rd in top performing methods for stage 1 models and 2nd for Stage 2 models. The models it was compared against included VoroMQA, ProQ2 and MULTICOM Constructs.

### F. Hybrid methods

Several methods have found success by combining existing methods in the field[31], [32]. ModFold6[30], [33] is one of the represented novel hybrid method for scoring predicted protein structures. The method utilizes both existing single-model methods and quasi-single methods in conjunction to assess quality scores. Both the sequence and the predicted structure are utilized in this method. The 3D structure is fed into 3 seperate single-model methods (CDA, SSA and ProQ2) as well as 3 Quasi-single methods (DBA, MF5s and MFcQs). The predicted residue scores from these methods were then fed into an artificial neural network using a sliding window of size 5. This neural networks output was single quality score for each residue. The paper presented a benchmark using area under the curve for a receiver operating characteristic. Using this metric the method outperformed ProQ3, Multicom-cluster and qSVMQA.

## III. Case study on Reinforcement learning Application

Reinforcement learning is one special type of AI that functions solely on numerical rewards derived from a particular environment. This technique is an extremely powerful tool in certain situations as it can function with sparse rewards and little training data. This is due to the fact that the model can generate its own training data through interaction with its environment. There has been much success recently in utilizing reinforcement learning and it has never been applied to the QA problem. As such, we will apply reinforcement learning in QA problem. Here we summarize the recent success it has seen in other fields. Q-Learning is the specific type of reinforcement learning that we would like to implement into our model.

Silver et al.[34] emphasises on using convolutional neural networks and reinforcement learning to teach an AI model how to play the game of Go by playing games against itself and then master the moves that it takes to win. The AlphaGo Zero model takes a picture of the board as input then calculates which move has the highest probablility of winning taking into account which move the opponent would make in response to Alpha’s move. The agent of the reinforcement learning is then handed the new state of the board and the reward for playing a move. The reward is whether or not that move helped AlphaGo Zero win the game. We would like to explore the reinforcement learning for QA method. Pathak et al.[35] emphasizes the importance of reinforcement learning. In many real-world applications there are very little extrinsic rewards to the agent exist or none at all. Also, often the agent cannot detect or construct the reward. Using this idea, this group produced a Super Mario world with no rewards. The only way the character could “learn” from the environment was from dying and teaching itself what caused it. There were no rewards provided for tasks such as killing enemies and avoiding fatal events. The character was able to discover these behaviors. This was the first known demonstration of learning to act in a relatively complex visual imagery directly from pixels without any extrinsic rewards. Q-Learning[36] is a form of reinforcement learning that provides a simple way to train a program to work the best in a controlled environment. It uses incremental methods which allows the program to train fast and requires small computational demands. It acts by continually re-evaluating an action to a reward whether it good or bad to update that particular action. This paper uses a theorem developed by Watkins in 1989. It showed that as long as all actions were repeatedly sampled and action values were represented discreetly the model could train to an optimum probability value of 1.

### A. Data formatting

The dataset we used is CASP dataset. The CASP Experiment releases targets weekly throughout the experiment with approximately 150 predicted models with varying levels of accuracy. For our experiment we processed the data into a JSON file using a program called Unicon3D[27]. This JSON file was organized by target with amino acid structures provided and 5 values of bond lengths and angles.

The initial data preparation was performed on CASP 10 data consisting of 36,083 models. Our first step was to filter these models by verifying that the predicted residue sequence matched that of the native protein and removing any nonmatching models or incomplete models. Next we create new representations of the models in the form of a position data file. The position data file represents a protein structure as an array of bond angles corresponding to each amino acid’s carbon alpha in a given peptide chain. Once we have the array of angle information we encode the amino acids and any other input features and join them to the angle information. At this point, we have a large numpy array representing one model. In order for our agent to learn, we must break this environment up into steps. We achieved this by slicing the large array into smaller arrays based on a window size. When combined, these slices will be one episode in the environment.

Support Vector Machines require data that has constant lengths. However, proteins vary greatly in amino acid chain length. One solution is to use primary component analysis (PCA) to give data constant length required by SVM. PCA takes given features then calculates the variance of each feature for a given data set and produces a fixed number of components that represent variance at each feature. PCA was used with 20 components and 5 features as the new dimensions for our matrix of features and amino acids from. The originally formatted CASP 10 json data was inverted so features were considered rows and amino acids were columns in order to get the amino acid variance of each feature. This data conversion is useful in taking 2D data like the bond angles vs amino acid into 1D data of the variance of bond angle that is represented by 20 components (roughly one component per known major amino acid.

### B. Reinforcement learning

The environment for our RL agent was created using the openAI gym interface. OpenAI gym is a platform for building, training and testing reinforcement learning models. To implement the environment we utilized the existing gym interface. Our environment consists of a numpy array of shape (modelName, gdt, img1, img2, …, imgN). Each row corresponds to the current model and contains the gdt score for that model and a sequence of images. The sequence of images are the previously sliced arrays and represent the bulk of the environment. The agent will receive three stacked images as an observation of the environment and choose whether to increase, decrease or leave the predicted gdt score alone. Each action is considered a step in the environment and upon taking that step the agent will receive the next three images for the current model. The end of an episode occurs when the end of the current model is reached and the reward used to train the agent is the negated absolute value of the difference between the gdt score and the predicted score. The agent will perform so many iterations per model. After each episode the environment is reset, this entails getting the next model for the agent and loading the first observation.

### C. Support Vector Machine

The Support Vector Machine used for protein prediction analysis was using the regression type to predict GDT scores given the bond angles and bond lengths. The original SVM was classification type and did not work on the data because the SVM could not classify the model as simply good or bad due to the output being a numerical GDT score. The regression type allowed for the use of a real number as the predicted GDT score. The data that was used in the SVM as input was first preprocessed in PCA in give each protein consistent length. The linear function was used as the kernel and The predicted GDT scores for each training model are then ran through the pipeline analysis to check the accuracy of the QA method. After running the SVM on the data, the true and predicted GDT scores were stored in seperate 2 dimensional arrays. That data was then sent through a for loop to compare to determine the Pearson’s Correlation Coefficients and GDT Loss for each target. Then that value for each target was averaged for the overall averages.

### D. Result of Reinforcement learning

All data used for training and testing was provided by CASP. CASP10 was used for training and CASP12 was used for testing. A total of 13944 models were used for training. Right now only 10 models were analyzed in the testing phase. The losses ranged from 0.14 to 0.39 with an average loss of 0.26. Correlation was not performed on this set of results due to the low number of models tested.

### E. Result of SVM

The data from CASP was used to train our SVM and then a homegrown pipeline analysis script was used to analyze GDT predictions. The correlation on CASP 10 and CASP 12 is – 0.005 and 0.016 respectively, and the loss is 0.092 and 0.198 respectively. The data produced by the svm can be shown graphically to illustrate why the correlation values are so low (**Figure 1**).

**Figure 1.**
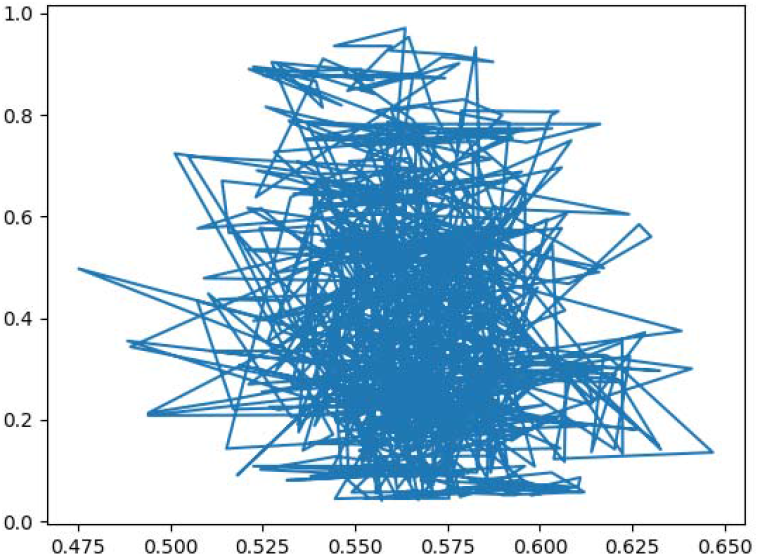
The x-axis is predicted GDT scores vs the y-axis which is actual GDT scores of proteins. The fitline is shown in blue.

The regression function is not fitting well with the data because all of the predicted GDT scores are clustered around 0.55. The major issue contributing to this is the loss of feature data such as angle and bond length through the PCA data downsizing. We tried varying different aspects of the SVM methods itself such as the kernel, C, and gamma variables as well as changing the output length of the PCA data in the hopes that we could retain more diverse data. Changing the way the 2D data of angles and amino acids is processed for the SVM would likely help to make a better spread and allow the SVM to fit a better function that produces higher correlation rates.

### F. Current Issues and learned lessons

There are currently many remaining issues with reinforcement learning and SVM in Bioinformatics tested by our methods. We select the following two points:

- Reinforcement learning: Currently the training loss is diverging. Several strategies have been attempted to correct this. Different learning rates have been tried. This parameter is currently set to 0.0001. Our hypothesis is that there is a problem with the ADAM optimizer that is used. We have added a clipnorm value of 1, and tried several different values of epsilon to no avail. More research is needed into why this is happening.
- SVM: The SVM is currently producing GDT scores that are low in correlation and high in loss. It is partially due to the formatting of the data that is put into the SVM. The data is preprocessed using PCA which causes information loss about the bond angle because the PCA formatted data presents a variance ratio of the angles rather than the angles themselves. The processed data is then used as input and the loss of information is compounded upon when the prediction is made using the PCA data. Another possible issue with the SVM is the setting of parameters, such as C and Gamma. We found that a linear fit for the kernel gave the best output. These issues with parameters however are likely on small issues compared to those posed by PCA.

The Reinforcement learning method is not showing the desired performance even with the few models that have been predicted. Additional features will be added including secondary structures. For the SVM method, we will try different parameters and more components for the PCA. We plan to try a different way of sizing the data to constant length so that we can run it through the SVM.

### G. Ethics in Bioinformatics

Right now, there are very few research on the Ethical issue in the Bioinformatics field. However, as we know, the success of human genome project helps all researchers to better understand our genome, and makes it possible for understanding all proteins in our body. The privacy issue is still a big problem, is that ok to release protein structure for a group of people? is the protein structure information considered as sensitive data for a person? How should we treat the world’s first genome-edited babies at the end of 2018 with the help of Crispr techniques? [37]. All of these questions should be carefully solved in the future research of applying AI in Bioinformatics.

### H. Future applications

AI while still having issues is overall beneficial for research in bioinformatics. First, AI allows for more accurate prognosis and diagnosis of structures because the computers can analyze data and have perfect calculations and deeply analyze the detail. These accuracies while may be very close to that of traditional approaches are still slightly stronger allowing confidence in the results. Second, AI is time and resource efficient. The results of an AI analysis come back much faster than a human could ever do. Also the researcher would be able to see other methods rather than spending time on analyzing images, structures and other modeling data. That could all be done by computer freeing up the researcher to carry out more conceptual analysis. Finally operating with AI is going to greatly cut costs and allow money to be used more efficiently. And the best part is that AI would not be replacing researchers but rather working in conjunction with them. The AI system would give suggestions and while researchers would make the final decision. AI will become more integrated with bioinformatics as the research broadens, and it will be interesting to see new benefits that arise as a result.

## IV. Conclusions

Artificial Intelligence is an exciting field when applied to research in bioinformatics. It offers solutions to issues in finding structures of proteins which is crucial to drug development and the understanding of biochemical effects. In this paper, we studied various In the general field of artificial intelligence, reinforcement learning is relatively undeveloped but very powerful. Such research proposes an additional foundation for the innovative process of protein folding predictions in general. A process for management of innovation in the research of bioinformatics will need to be worked out. Specific algorithms can be developed for research in protein folding predictions. In future, we need to apply various AI techniques and explore different reinforcement learning techniques to improve the performance of available AI methods.

## Acknowledgment

We would like to thank the support from Pacific Lutheran University.

